# Chronic IL-6 overproduction induces a tolerogenic response in aged mice after peripheral nerve injury

**DOI:** 10.1101/2024.03.13.584805

**Authors:** Gemma Manich, Ruggero Barbanti, Marta Peris, Nàdia Villacampa, Beatriz Almolda, Berta González, Bernardo Castellano

## Abstract

**Highlights/Main points:**

- **Astrocyte-targeted IL-6 overproduction during aging increases basal microglial acivation in the facial nucleus.**
- **During aging, chronic IL-6 overproduction does not modify microglial response after peripheral nerve injury but increases T-lymphocyte infiltration.**
- **Chronic IL-6 overproduction in aged mice does not modify facial motor neuron survival after facial nerve axotomy.**

Interleukin-6 (IL-6) is the main cytokine controlling neuroinflammation and microglial activation during aging, and the increase of IL-6 levels correlate well with chronic neuroinflammation and age-related neurodegenerative diseases. Despite the relevance of IL-6 in these conditions, the effect of this cytokine in microglia activation and neuroinflammation upon CNS injuries during aging has been scarcely explored. Previous results from our group showed that adult and aged transgenic mice with astrocyte-targeted overproduction of IL-6 (GFAP-IL6Tg) presented features of a primed microglial phenotype in basal conditions compared to wild-type (WT) mice, and that after CNS lesions, microglia showed and exacerbated response associated with increased neuronal death in adult mice. In this work, we aimed to study whether chronic IL-6 overproduction influenced microglia response to CNS injury during aging. With this aim, we performed facial nerve axotomy (FNA) in aged 21-23-month-old WT and GFAP-IL6Tg animals, and analysed facial motor neuron (FMN) survival, glial reactivity, antigen presentation, and lymphocyte infiltration both at basal conditions (non-lesioned) and after FNA. Our results showed that non-lesioned aged transgenic mice presented higher Iba1, CD11b, and CD68 levels than aged WT mice, indicative of a priming effect in the aged facial nucleus. After FNA, we detected similar levels of microglial and astroglia activation, but a remarkable increase in T-lymphocyte infiltration in GFAP-IL6Tg axotomized group. Despite slight differences in the neuroinflammatory response, aged GFAP-IL6Tg animals showed a similar rate of FMN death compared to aged WT mice. Overall, our work shows that transgenic IL-6 overproduction induces a primed microglial phenotype under basal conditions in aged animals, with a reduced fold-increase in the microglial response after FNA compared to aged WT mice and similar lesion outcomes, suggestive of a tolerant microglial phenotype. This study suggests a tolerant effect of chronic IL-6 overproduction in microglia during aging in basal conditions and after CNS lesions.

## INTRODUCTION

Physiological aging in the CNS is associated to a chronic low neuroinflammation grade produced in the tissue, also named ‘inflammaging’ (Calabrese et al., 2018; Franceschi et al. 2017), resulting from an imbalance caused by increased levels of pro-inflammatory cytokines and decreased levels of anti-inflammatory cytokines (Antignano et al., 2023; Udeochu et al. 2016). ‘Inflammaging’ promotes a shift in microglia to a pro-inflammatory phenotype in aged mice (von Bernhardi et al. 2015), even in newly generated one (Elmore et al. 2018, O’Neil et al. 2018; Stojiljkovic et al., 2022). This specific aged microglial phenotype is characterized by soma hypertrophy, shorter and thicker ramifications, reduced process motility and chemotactic capacity (Wong et al., 2013; Hefendehl et al. 2014; Rawji et al. 2018; Spittau, 2017), impaired phagocytosis (Safaiyan et al. 2016; Rawji et al. 2018), and altered transcriptome compared to homeostatic microglia (Grabert et al., 2016; Hammond, 2019; Hickman et al. 2013; Holtman et al., 2015; Marschallinger et al., 2020; Silvin et al., 2022).

One of the main contributors to ‘inflammaging’ is the pro-inflammatory cytokine interleukin (IL)-6. This cytokine has been strongly and positively correlated to decreased life expectancy in old individuals, and to age-related decrease in motor and cognitive performance (Richwine et al. 2005; Guest et al. 2014; McCarrey et al. 2014; Varadhan et al. 2014; Niraula et al., 2017). Some works showed that IL-6 is upregulated in the brain of aged mice not only under homeostatic circumstances, but also after both central and peripheral insults, produced, for example, by LPS administration (Ye and Johnson 2001; Godbout and Johnson 2004; Godbout et al. 2005; Huang et al., 2008).

In the CNS, this cytokine is mainly produced by microglia upon inflammatory stimulus, and since it is one of the first cytokines to be released in inflammation, it regulates the pro-inflammatory phase of this process. IL-6 controls neuroinflammation through two signaling pathways: 1) the classical pathway, through the IL-6 receptor found in microglia, among other cell types, in the CNS, and 2) the trans-signaling pathway, activated by means of the soluble IL-6 receptor, which can signal in most CNS cell types (for reviews see Erta et al., 2012; Rothaug et al., 2016; Scheller et al., 2011; Spooren et al., 2011; West et al., 2019). Despite its overspread signal in the CNS, IL-6 produces the most remarkable effects over microglia itself, its main producers, leading to a pro-inflammatory loop.

The effects of IL-6 on microglia include microgliosis, alterations in morphology-shortening of ramifications, less branching complexity and larger somata-, upregulation of MHC microglial expression, increased complement pathway activation and enhancement of the microglial phagocytic phenotype, as well as a metabolic shift to glycolytic pathways, among other effects (Barnum et al., 1996; Campbell et al., 1993; Fattori et al., 1995; Gyengesi et al., 2018; Petkovic et al., 2016; Sanchez-Molina et al., 2021; Shafer et al., 2002; West et al., 2019, 2022a). Chronic astrocyte-targeted overproduction of IL-6 in transgenic mice (GFAP-IL6Tg) induced changes in the microglial transcriptome genes related to phagocytosis engulfment and processing of cell remnants, regulation of iron metabolism and angiogenesis. Remarkably, the GFAP-IL6Tg microglial transcriptome showed common microglial gene cluster expression compared to LPS-administered mice (West et al., 2022b). As we and others have observed, IL-6 induces a priming effect on microglia (Garner et al. 2018; Recasens et al. 2021; Sanchez-Molina et al. 2021), and after a lesion, when this cytokine is chronically expressed in adult mice, the microglial phagocytic capacity and activation is modified, up to the point to influence the lesion outcome (Recasens et al., 2021; Manich et al., 2020; Petkovic et al., 2016). Specifically, after a peripheral nerve injury, adult GFAP-IL6Tg mice showed higher facial motor neuron death compared to injured wild-type (WT) mice, tightly related to a dysfunctional microglia activation (Almolda et al. 2014).

In this work, we hypothesize that chronic IL-6 overproduction during aging will result in a sustained pro-inflammatory situation and microglial priming that will alter the neuroinflammatory response to a CNS lesion. Our aim is to study the effect of chronic IL-6 overproduction in neuroinflammation and neuronal survival after facial nerve axotomy (FNA) during aging.

## MATERIALS AND METHODS

### Animals

In the present work, 28 GFAP-IL6Tg animals (Campbell, Abraham et al. 1993), and 26 C57BL/6 wild-type (WT) littermates, aged 21-23 months old, from both sexes were used. All mice were housed under a 12h light/dark cycle, with food and water *ad libitum*. All experimental animal work was conducted according to Spanish regulations (Ley 32/2007, Real Decreto 1201/2005, Ley 9/2003 and Real Decreto 178/2004) in agreement with European Union directives (86/609/CEE, 91/628/CEE and 92/65/CEE) and was approved by the Ethical Committee of the Autonomous University of Barcelona. Every effort was made to minimize the number of animals used to produce reliable scientific data, as well as animal suffering and pain.

### Facial nerve axotomy and experimental groups

GFAP-IL6Tg and WT mice were anaesthetized with a solution of ketamine (80 mg/kg) and xylazine (20 mg/kg) intraperitoneally injected at dose of 0.01 ml/g. The skin behind the right ear was shaved and cleansed with 70% ethanol. A small incision was made in the skin and both the trapezius and the anterior digastric muscles were gently separated to expose the right facial nerve. One millimeter of the facial nerve main branch was resected at the level of the stylomastoid foramen. Following the surgery, the skin was sutured with 5-0 nylon. Corneal dehydration was prevented by application of Lacri-lube® eye ointment. After anesthesia recovery, the complete whisker paralysis was assessed to ensure that the complete facial nerve resection was achieved. Axotomized animals were distributed in different experimental groups and euthanized at 14- and 21-days post-injury (dpi) according to the appropriate experimental procedure. A group of non-lesioned (NL) WT and transgenic animals was also included in the study and used as control.

### Tissue processing for histological analysis

Animals were anaesthetized intraperitoneally with the same ketamine/xylazine solution but used at dose of 0.015 ml/ g and perfused intracardially with 4% paraformaldehyde in 0.1 M phosphate buffer (pH 7.4). Brains were removed and post-fixed in the same fixative for 4h at 4°C and, after rinsing in phosphate buffer (pH=7.4), they were cryopreserved for 48h at 4°C in a 30% sucrose solution and subsequently frozen with 2-methylbutane solution (Sigma-Aldrich). Parallel free-floating coronal sections (30-µm-thick) of the brainstem containing the facial nucleus (FN) were obtained using a CM3050s Leica cryostat and sections stored at - 20°C in Olmos antifreeze solution until their use.

### Single immunohistochemistry

Free-floating cryostat sections were processed for immunohistochemistry against Iba-1, CD11b, CD68, MHC-II, Pu.1, CD3 and GFAP. Briefly, for immunohistochemistry, sections were washed with Tris-buffered saline (TBS, pH 7.4) and inhibited for endogenous peroxidase by incubating samples with a solution of 2% H_2_O_2_ in 70% methanol for 10 min. For the detection of Pu.1 transcription factor, an antigen retrieval step was previously performed, by incubating sections for 20 min at 98°C in 10mM sodium citrate buffer (pH 8.0). Afterwards, sections were incubated for 1h in blocking solution containing 10% foetal calf serum and 3% bovine serum albumin (BSA) in TBS with 1% Triton X-100 (TBS-T). Then, sections were incubated with rabbit anti-Iba1 (1:1000; GeneTex), rat anti-CD11b (1:1000; MCA74G, AbD Serotec), rat anti-CD68 (1:1000; MCA1957, AbD Serotec), rat anti-MHC-II (1:25, rat hybridoma TIB-120, ATCC), rabbit anti-Pu.1 (1:200; 2258, Cell Signalling), hamster anti-CD3 (1:500; MCA2690, AbD Serotec) or rabbit anti-GFAP (1:2400, Z0334, Dako) diluted in the blocking solution overnight at 4°C plus one additional hour at room temperature (RT). Negative controls were performed using sections incubated in media lacking the primary antibody, and spleen sections were used as positive control. After washing three times with TBS-T, sections were incubated at RT for 1h with biotinylated anti-rabbit (1:500; BA-1000; Vector Laboratories, Inc; Burlingame, CA), biotinylated anti-rat (1:500; BA-4001; Vector Laboratories) or biotinylated anti-hamster (1:500; BA-9001; Vector Laboratories) secondary antibodies diluted in the blocking solution. After 1h at RT in horseradish streptavidin-peroxidase (1:500; SA-5004; Vector Laboratories), the final reaction was visualized by incubating the sections with a DAB kit (SK-4100; Vector Laboratories) following the manufacturer’s instructions. Finally, sections were mounted on gelatin-coated slides, counterstained with toluidine blue, dehydrated in graded alcohols and, after xylene treatment, coverslipped with DPX. Sections were analyzed and photographed with a DXM 1200F Nikon digital camera joined to a Nikon Eclipse 80i microscope.

### Double immunohistochemistry

Free-floating cryostat sections were processed for double immunohistochemistry against MHC-II and laminin. Briefly, sections were washed with TBS and were incubated for 1h in blocking solution. Then, sections were incubated with rat anti-MHC-II (1:25, rat hybridoma TIB-120, ATCC) diluted in the blocking solution overnight at 4°C plus one additional hour at RT. After washing three times with TBS-T, sections were incubated with AlexaFluor 555 anti-rat antibody (1:500; A21434; ThermoFischer Scientific) diluted in blocking solution for 1h at RT and washed with TBS-T. Then, sections were incubated again for 1h with blocking solution, and then with the rabbit anti-laminin primary antibody (1:500; L9393; Sigma-Aldrich) diluted in the blocking solution overnight at 4°C plus one additional hour at RT. After washing sections with TBS-T, sections were incubated with AlexaFluor 488 anti-rabbit antibody (1:500; A21206; ThermoFischer Scientific) diluted in blocking solution for 1h at RT. Sections were washed with TBS and then TB; and counterstained with DAPI (1:10,000; D9542; Sigma-Aldrich) diluted in TB for 5 min. Finally, sections were mounted on gelatin-coated slides and coverslipped with Fluoromount-G Mounting Medium (00-4958-02; ThermoFischer Scientific). Negative controls were performed using sections incubated in media lacking the primary antibody, and spleen sections were used as positive control. Sections were analyzed and photographed with a Zeiss LSM 700 confocal microscope.

### Neuronal cell counting of the facial nucleus

Consecutive cryostat sections (30-μm-thick) from 21 dpi animals containing the ipsilateral (axotomized) and contralateral (non-axotomized) FN were mounted on gelatin-coated slides and stained with a 0.1% toluidine blue solution diluted in Wallpole buffer (0.05 M, pH 4.5). After staining, sections were dehydrated in a sequence of graded alcohols, N-butyl alcohol and xylene and coverslipped with DPX.

The contralateral and the ipsilateral side of every section through the entire FN was examined and photographed using a DXM1200F Nikon digital camera joined to a brightfield Nikon Eclipse 80i microscope. The photographs were analyzed with the digital AnalySIS® software (Soft Imaging System). In addition to the total number of neurons, the maximum and the minimum diameters of neuronal profiles in the FN were recorded to obtain the mean diameter. To compensate for double neuronal count in adjacent sections, the Abercrombie’s correction factor was applied as previously reported (Almolda et al. 2014, Villacampa et al. 2015).

### Brightfield microscopy quantification

To study microglial and astroglial reactivity, quantitative analysis was performed on sections individually immunolabeled for Iba-1, CD11b, CD68, MHC-II, and GFAP. Between three to four GFAP-IL6Tg and their corresponding WT littermates were analyzed at each timepoint (NL, 14 and 21 dpi). For each animal, at least three representative sections from the brainstem containing the central part of the ipsilateral or contralateral FN were photographed at 10X magnification. For each marker, the percentage of the area covered by the immunolabeling (% Area) as well as the intensity of the immunoreaction, measured as the Mean Grey Value and expressed in arbitrary units, were analyzed using the AnalySIS ® software (Acarin et al. 1997).

For determination of microglia cell density, Pu.1 quantification was performed in ipsilateral and contralateral FN of at least three WT and three GFAP-IL6Tg of each timepoint (NL, 14 and 21 dpi). The FN areas of at least three representative sections for each animal were captured at 10x, and images were then analyzed by counting the total number of Pu.1+ cells in each section using the “Automatic nuclei counter ICTN” plugin of the NIH ImageJ software (version 1.51).

Quantification of microglial clusters, i.e., groups of a minimum of 4 or more nuclei of microglial cells, was performed at 14 and 21 dpi on sections stained for both Iba1 and Pu.1, in a minimum of three WT and five GFAP-IL6Tg animals. At least three representative sections from the brainstem containing the central part of the FN from each animal were observed in the microscope and counted manually. The number of clusters was expressed as Iba-1-positive or Pu.1-positive clusters/section.

To determine T-lymphocyte infiltration in the FN, CD3-positive cells were counted in three WT and three GFAP-IL6Tg animals of each timepoint (NL, 14 and 21 dpi). At least six representative sections containing the ipsilateral or contralateral FN for each animal were captured at 10x magnification. All CD3-positive cells in each microphotograph were counted using the “Cell counter” plug-in from ImageJ software (National Institutes of Health). Results were expressed as CD3-positive cells/section.

### Tissue processing for protein quantification

Animals were anaesthetized intraperitoneally with ketamine/xylazine at dose of 0.015 ml/g and perfused intracardially for 1 min with 0.1 DPBS (D8537, ThermoFisher Scientific). Brains were quickly removed, and each FN was dissected from two 0.5-mm-thick coronal slices of the brainstem obtained using a Mouse Brain Matrix (Zivic Instruments) including areas from bregma −5.52 to −6.72 mm approximately, and by cutting the dorsal inferior half of the brainstem with sterile knives. The tissue was immediately snap frozen in liquid nitrogen and stored at −80°C. Total protein was extracted by sample solubilization using lysis buffer containing 25mM HEPES, 2% Igepal, 5mM MgCl2, 1.3 mM EDTA, 1 mM EGTA, 0.1 M PMSF and protease (1:100, P8340; Sigma-Aldrich), and phosphatase inhibitor cocktails (1:100, P0044; Sigma-Aldrich) for 2 hours at 4°C. Then, samples were centrifuged at 13000 rpm for 5 min at 4°C and the supernatants collected. Total protein concentration was assessed with a commercial Pierce BCA Protein Assay kit (23225; ThermoFisher Scientific) according to manufacturer’s protocol. Protein lysates were aliquot and stored at −80°C until used for protein microarray analysis. The FN of each animal was analysed separately.

### Quantification of cytokines by Multiplex analysis

IL-6, IL-10 and TNF-α were quantified in the FN of NL and at 21dpi aged GFAP-IL6Tg and WT mice (n= 4-6/experimental group) using a Milliplex MAP Mouse High Sensitivity kit (#MHSTCMAG-70K; Merck Millipore), following the manufacturer’s instructions. Briefly, 25µL of standards and each FN extract (2.0 µg/µL total protein) were added to wells, containing 25µL of custom fluorescent beads and 25µL of matrix solution, and incubated overnight at 4°C in a plate-shaker (750rpm). After washing twice with wash buffer, the plate was incubated with 25µL of detection antibodies for 30 min at RT, and with 25µL of Streptavidin-Phycoerythrin for 30 min at RT in a plate-shaker (750rpm). Finally, the plate was washed twice using wash buffer and 150µL of Drive fluid was added. Luminex MAGPIX device with the xPONENT 4.2 software was used to read the plate. Data were analysed using the Milliplex Analyst 5.1 software and expressed as pg/mL of protein.

### Quantification of soluble IL-6 receptor (sIL-6R)

Determination of sIL-6R was performed in the FN of NL and 21 dpi of aged WT and GFAP-IL6Tg (n=3-6/experimental group) using a precoated mouse ELISA kit (EMIL6RA, Fisher Scientific) and following the manufacturer’s instructions. Briefly, 100µL of standards (6.14 - 1500pg/mL) or FN extract (0.5-1.0 µg/µL total protein) were added to wells and incubated overnight at 4°C with agitation. After several washes, the biotin-conjugated antibody was added, and after washing again, streptavidin was incubated at RT, and washed. Then, the TMB substrate was incubated for 30 min, the stop solution was added and results were read at 450 nm in the microplate reader VarioskanTM Lux (Thermo Fisher Scientific).

### Statistical analysis

Statistics were performed using Graph Pad Prism® software version 6 (Graph Pad Software Inc.) and results were expressed as mean ± standard error of the mean (SEM). To detect differences in basal conditions, a standard two-tailed unpaired Student’s t-test was used for comparison between NL GFAP-IL6Tg and NL WT mice. Two-way analysis of variance (ANOVA) was used for comparisons between groups from NL, 14 and 21 dpi, and post-hoc Tukey’s or Sidak’s Multiple Comparison Test was applied to compare among groups.

## RESULTS

### Determination of IL-6 expression in the FN during basal conditions and after FNA in aged WT and GFAP-IL6Tg mice

First, we evaluated the levels of IL-6 in the FN of the non-lesioned (NL) and lesioned (21dpi) aged WT and GFAP-IL6Tg using the Luminex microarray of proteins. Our results showed that IL-6 levels are higher in the FN of aged GFAP-IL6Tg, both in basal conditions and after injury, compared to WT (Figure 1).

**Figure 1.**
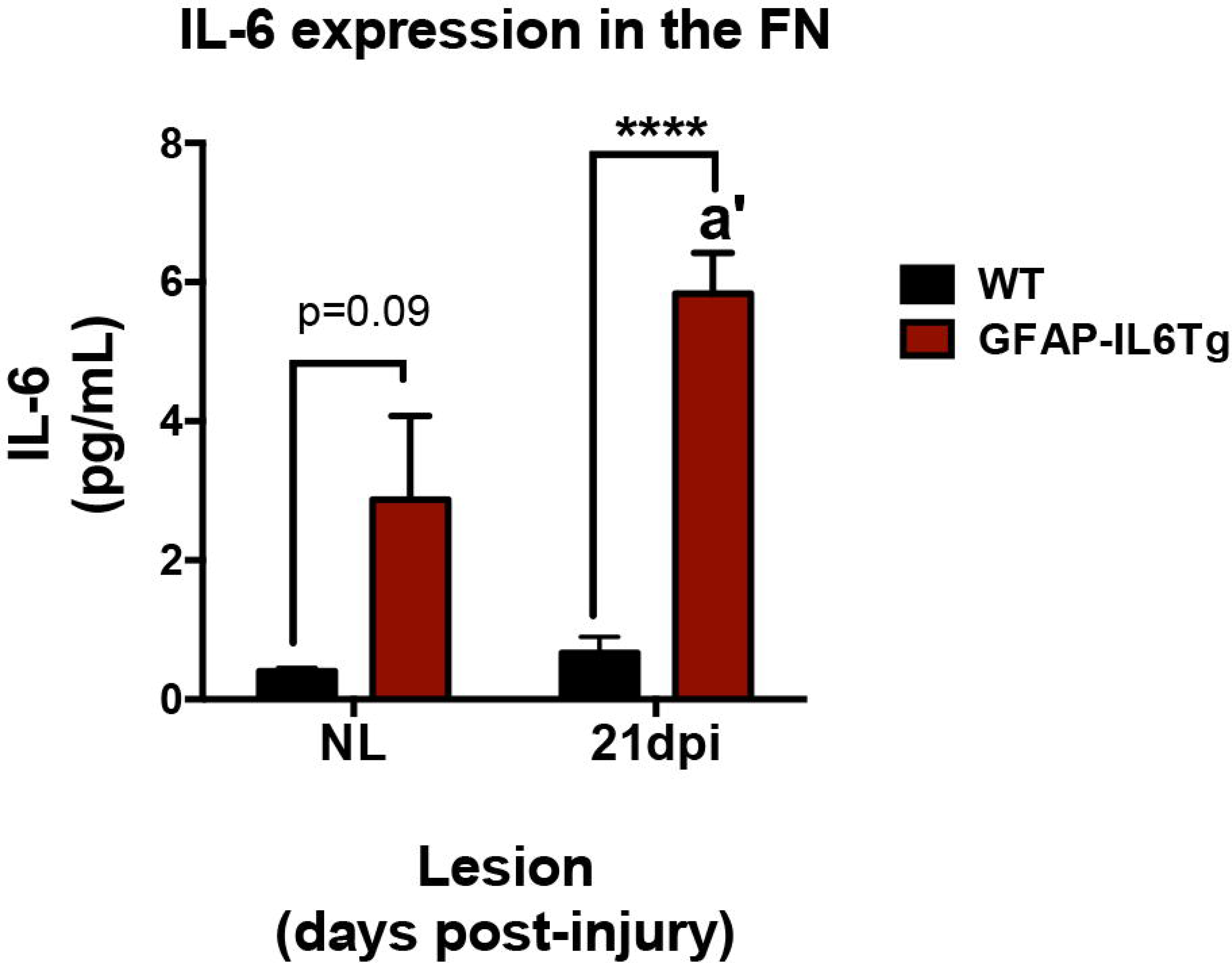
IL-6 expression in the FN of aged WT and GFAP-IL6Tg mice during basal conditions and after FNA. Quantification of IL-6 expression was performed in the NL (non-lesioned) and lesioned (21 dpi) FN of aged WT and aged GFAP-IL6Tg. IL-6 expression was increased in the FN of aged GFAP-IL6Tg in NL and lesioned conditions when compared to the corresponding aged WT (2-way ANOVA, genotype p<0.0001, lesion p<0.05, interaction p<0.05; post-hoc Tukey’s, a’ p<0.05 compared to NL, **** p<0.0001).

### Microglial characterization in basal conditions and after FNA

a) *Microglia morphology and distribution:*

The evaluation of morphology, distribution, and activation of microglia in the FN of aged mice in basal conditions and after FNA, was performed through Iba-1 and Pu.1 immunostains (Figure 2). No significant differences were observed when comparing the FN of NL animals and the contralateral FN of axotomized mice (data not shown), either in WT or GFAP-IL6Tg groups, and consequently in the entire study we only used the contralateral FN for statistical analysis.

**Figure 2.**
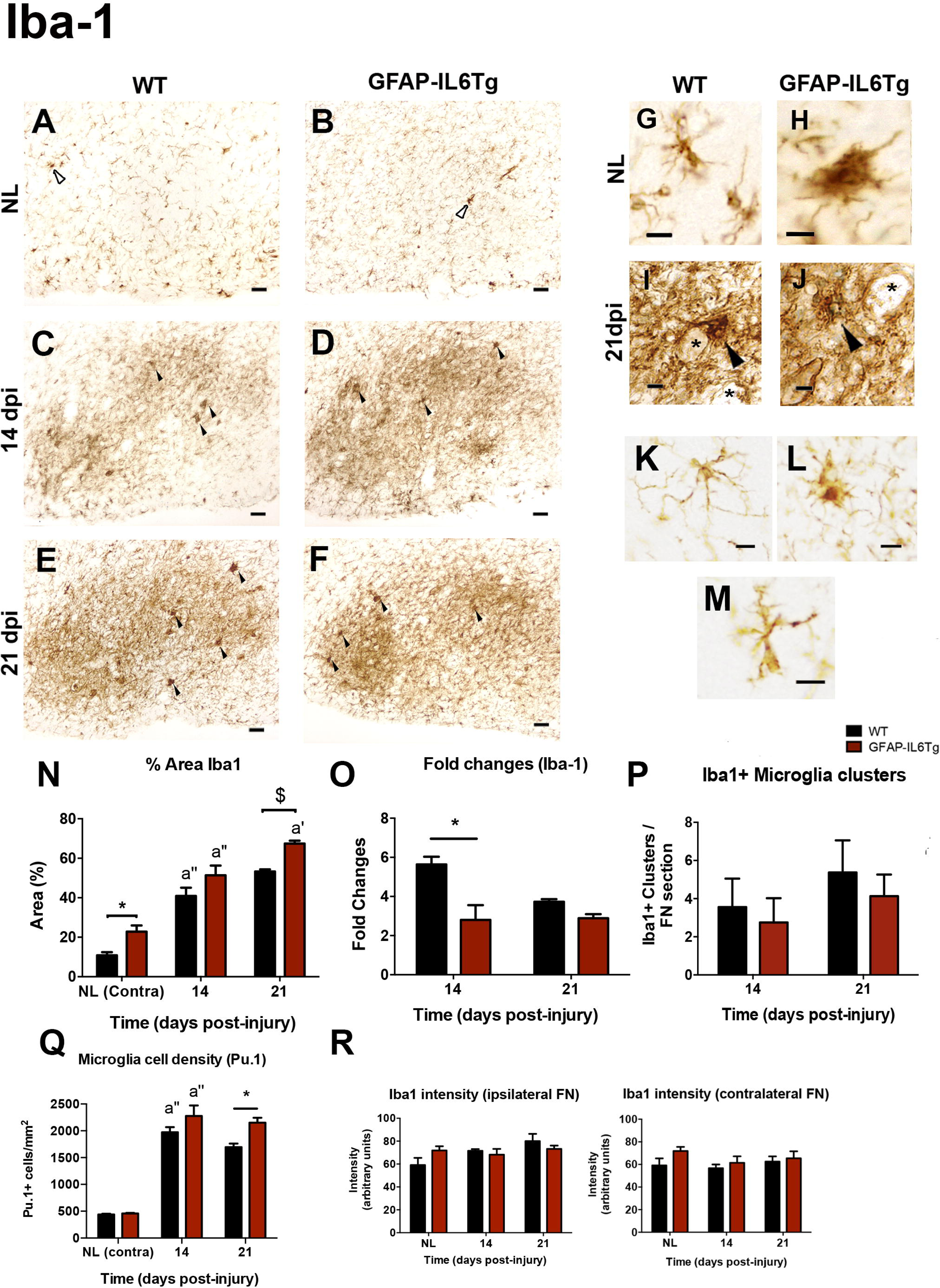
Microglial reactivity in basal conditions and after a FNA in aged WT and GFAP-IL6Tg mice. **(A-F)** Representative photomicrographs from Iba-1 immunostaining in 21-23 months-old WT and GFAP-IL6Tg mice, in both basal conditions and after peripheral nerve injury. In the NL FN, bushy microglia is observed (empty arrowheads), and after lesion microglial clusters are formed (full arrowheads) **(G-J)** Magnification of brain parenchyma showing microglial morphology. Note the altered microglial morphology in basal conditions in aged WT mice (G) and bushy microglia in GFAP-IL6Tg (H). After the axotomy, enhanced microglial reactivity and microglial clusters (arrowheads) next to FMN (asterisks) are observed (I, J). (**K-M)** Microglial morphologies observed in the aged NL FN of WT and GFAP-IL6Tg mice. Microglia with heterogeneous morphology were observed: microglia with extended and abundant ramifications (K), microglia with a bushy appearance (L) and microglia with short branches and enlarged soma (M). **(N)** Quantification of the % Area of Iba-1 immunoreactivity in WT and GFAP-IL6Tg in the ipsilateral FN. Note that basal % Area in contralateral hemispheres (NL Contra) are increased respect to aged WT mice (2-way ANOVA; *p<0.05). Higher levels of % Area are observed at 14 dpi in WT and at 14 and 21 dpi in GFAP-IL6Tg respect to the previous time-point (2-way ANOVA, time p<0.0001, genotype p<0.001; post-hoc Tukey’s, a’ p<0.05, a’’ p<0.0001). At 21 dpi, a tendency to increase in the % Area of Iba-1 is shown in GFAP-IL6Tg respect to WT mice ($ p=0.089). **(O)** Fold changes of Iba-1 microglial reactivity observed in lesioned compared to the respective NL FN in both aged WT and GFAP-IL6Tg. At 14 dpi, aged GFAP-IL6Tg show a lower increase respect to the NL FN when compared the increase observed in lesioned aged WT mice (2-way ANOVA, genotype p<0.01, post-hoc Sidak’s, *p<0.05). **(P)** Number of microglia clusters per FN sections were also counted at 14 and 21 dpi in both mouse lines, and no differences were found **(Q)** Microglial cell density was determined by counting Pu.1+ cells/mm^2^. In basal conditions WT and GFAP-IL6Tg presented similar microglia numbers, and at 14 dpi the number of microglia increased respect to NL (2-way ANOVA, time p<0.0001 genotype p<0.01; post-hoc Tukey’s, a’’ p<0.0001). At 21 dpi, GFAP-IL6Tg showed higher microglia numbers compared to WT mice (2-way ANOVA; *p<0.05) **(R)** Intensity of Iba-1 staining in the ipsilateral and contralateral FN of aged WT and GFAP-IL6Tg does not reflect statistically significant differences. Scale bar (A-F) = 50 µm. Scale bar (G-M) =10 μm.

In comparison to the typical ramified microglia of the FN with homogeneous parenchymal distribution described in adult animals (Carvalho-Paulo et al., 2021), microglia in aged NL animals showed a heterogeneous morphology in both WT and GFAP-IL6Tg (Figure 2A, B, G, H). Three morphological different cell types were identified in aged animals: 1) most microglial cells were characterized by branches with few ramifications, enlarged soma and short and coarse branches (Figure 2G, M); 2) microglial cells with extended branches and abundant ramifications –specially seen with CD11b immunostaining, as detailed in the following section– (Figure 2K), and 3) microglia cells with bushy appearance, occasionally clustered (empty arrowheads, Figure 2A, B, H, L). All those features were compatible with a high activated morphology and were observed in both WT and GFAP-IL6Tg.

**Figure 3.**
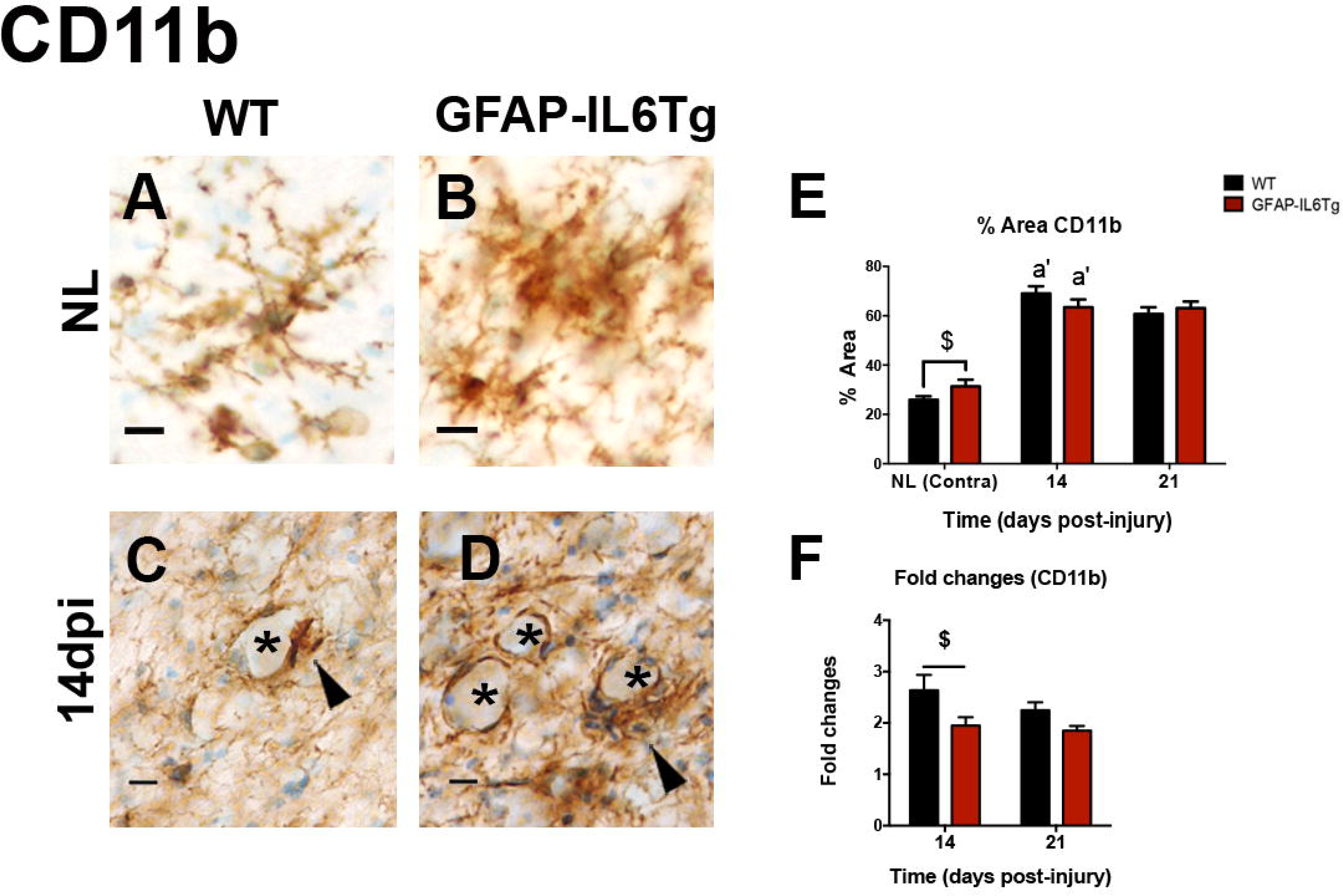
Microglial immunostaining with CD11b in basal conditions and after a FNA in aged WT and GFAP-IL6Tg mice. **(A-D)** Representative photomicrographs from CD11b staining show hyper ramified microglial morphology (A), altered microglial morphology and high immunostaining or bushy appearance (B) in non-lesioned (NL) aged WT mice (A) and in aged GFAP-IL6Tg (B). After injury, microglia wrapped FMN (**\***) and microglial clusters were formed (arrows) around FMN (C, D). **(E)** Quantification of the % Area with CD11b immunoreactivity in WT and GFAP-IL6Tg. Basal % Area in contralateral FN is increased in aged GFAP-IL6Tg respect to aged WT mice (Student’s t-test, $ p=0.07). Higher levels of % Area are observed at 14 dpi in both WT and GFAP-IL6Tg respect to basal conditions (2-way ANOVA, time p<0.0001; a’ p<0.0001). **(F)** Fold changes of CD11b immunostaining observed in lesioned compared to the respective NL FN in both aged WT and GFAP-IL6Tg. At 14 dpi, aged GFAP-IL6Tg show a lower increase respect to the NL FN when compared the increase observed in lesioned aged WT mice (2-way ANOVA, genotype p<0.01, post-hoc Sidak’s, $ p=0.09). Scale bar: (A-D) = 10 μm.

In accordance with these qualitative observations, the quantitative analysis of Iba-1 showed a significant increase in the percentage of stained area in the contralateral FN of aged GFAP-IL6Tg animals (Figure 2N), but without significant differences in the immunostaining intensity (Figure 2O, P. This increase in percentage of Iba1 was not due to modifications in the cell number, as according to Pu.1, no differences in the number of microglial cells between NL aged WT and NL aged GFAP-IL6Tg animals were found (Figure 2Q).

After FNA, an increase in microglial reactivity was found at 14 and 21 dpi in both WT and GFAP-IL6Tg mice. Reactive microglia showed thicker branches and higher Iba-1 expression (Figure 2C, D, N), and clustering (arrowheads, Figure 2C-F, I, J). Although the Iba-1 expression pattern was similar in the axotomized FN of both aged WT and GFAP-IL6Tg mice, the percentage of area covered by Iba-1 was increased at 21 dpi in transgenic animals (Figure 2N). Considering that the basal level of Iba1 was higher in NL conditions, as specified before, we calculated the fold-change increase of Iba-1 in comparison to their corresponding NL. These results demonstrated that, although the total amount of Iba1 was higher in transgenic animals, the increase in Iba1 after lesion was lower in GFAP-IL6Tg than in WT at 14 dpi (Figure 2R). In parallel to changes in expression, a higher number of microglial cells (Pu.1+) was found at 21dpi in GFAP-IL6Tg animals compared to WT (Figure 2Q). However, no differences between genotypes were observed in the microglial cluster formation (Figure 2 I, J, S). Moreover, at these specific time-points microglial cells and their processes completely wrapped the cell body of facial motor neurons (FMN). This microglial wrapping was found in both aged WT and aged GFAP-IL6Tg without detectable differences among genotypes (asterisks in Figure 2I, J).

b) *CD11b expression:*

In previous work from our group, we reported a dysregulated pattern of CD11b expression in microglia after FNA in lesioned GFAP-IL6Tg (Almolda, Villacampa et al. 2014). This observation correlated with decreased microglial attachment to FMN, and since CD11b exerts functions as integrin, CD11b deficiency could be the cause of increased facial motor neuron death observed in the adult transgenic animals (Almolda et al. 2014). Due to the modifications in Iba1 and Pu.1 in aged transgenic microglia, we decided to explore CD11b expression in microglia after FNA in aged WT and aged GFAP-IL6Tg.

In the NL FN of aged WT and GFAP-IL6Tg we identified the same microglial morphologies detected with Iba-1 (Figure 3A, B). Quantification of CD11b immunostaining showed that contralateral NL, FN, CD11b expression (calculated as the percentage of positive stained area) were similar in the contralateral FN of aged GFAP-IL6Tg compared to aged WT animals (Figure 3E). After FNA, in both WT and GFAP-IL6Tg mice, microglial reactivity increased (Fig. 3E), formed clusters and wrapped FMN somas in both aged, lesioned WT and GFAP-IL6Tg (Figure 3C, D; arrowheads and asterisks, respectively). As similarly described for Iba-1 immunostaining, aged GFAP-IL6Tg animals showed a reduced CD11b fold-change increase in relation to the NL FN at 14 dpi compared to lesioned WT animals (Figure 3F).

c) *Phagocytic capacity:*

In comparison to the slight CD68 lysosomal staining observed in the soma of microglial-like cells in NL aged WT mice (Figure 4A and G), CD68 staining in aged NL GFAP-IL6Tg mice was spread throughout all the cell body (Figure 4B and H). In agreement, the quantitative analysis showed an increase in the percentage of the area stained with CD68 in transgenic animals (Figure 4N), without differences in the intensity of the staining (Figure 4M).

**Figure 4.**
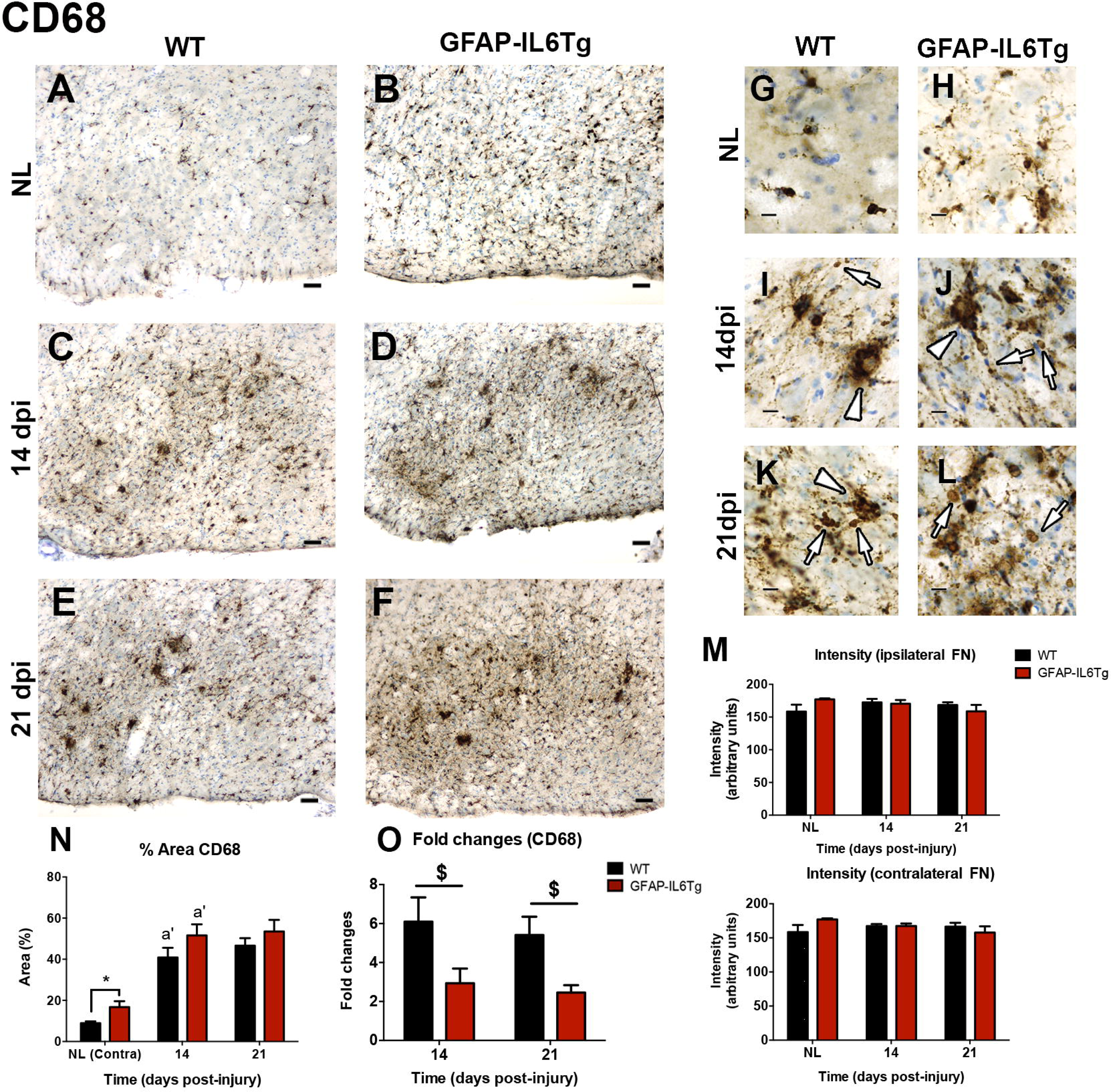
CD68 expression in basal conditions and after a FNA in aged WT and GFAP-IL6Tg mice. **(A-F)** Representative photomicrographs from 21-23 months-old WT and GFAP-IL6Tg mice in non-lesioned mice (NL) and at 14 and 21 days after a peripheral nerve injury. **(G-L)** Magnification of the FN showed microglial CD68 strong staining localized in the cell body in NL aged WT mice (G), while in NL GFAP-IL6Tg microglia phenotype, CD68 punctate staining seems to be spread throughout the cell body (H). After injury, CD68 lysosomal expression increased, was spread through cell bodies (arrows in I-L) and increased in size from 14 to 21 dpi in both mouse lines (empty arrows in I-L). CD68 expression was also concentrated in microglial clusters (empty arrowheads in I-L) in both 14 and 21 dpi. **(M)** Intensity of CD68 staining in the ipsilateral and contralateral FN of old WT and GFAP-IL6Tg at all timepoints remains constant. **(N)** Quantification of the % Area of CD68 immunoreactivity in WT and GFAP-IL6Tg shows that in basal conditions, % Area of CD68 was increased in aged GFAP-IL6Tg respect to aged WT mice (Student’s t-test; * p<0.05). At 14 dpi, both WT and GFAP-IL6Tg were increased respect to basal conditions (2-way ANOVA, time p<0.0001, genotype p<0.05; post-hoc Tukey’s, a’ p<0.0001). **(O)** Fold changes of CD68 immunostaining observed in lesioned compared to the respective NL FN of both aged WT and GFAP-IL6Tg. At 14 and 21 dpi, aged GFAP-IL6Tg tended to show a lower increase respect to their NL FN when compared the increase observed in lesioned aged WT mice (2-way ANOVA, genotype p<0.01, post-hoc Sidak’s, $ p=0.055). Scale bar (A-F) = 50 µm. Scale bar (G-L) =10 μm.

Upon FNA, at 14 dpi, CD68 expression increased in aged WT and GFAP-IL6Tg mice, and the immunostaining was found spread along reactive microglial cells in both groups (Figure 4C, D, I, J and N). At 21 dpi, CD68 expression levels remained unaltered in both axotomized WT and GFAP-IL6Tg respect to 14 dpi (Figure 4E, F and N). After the lesion, no significant differences between WT and GFAP-IL6Tg animals were found in neither the percentage of CD68+ area nor the intensity of the staining (Figure 4N and M). Again, when we calculated the fold-change increase of CD68 immunostaining at 14 and 21 dpi in relation to the values observed in their respective NL, we observed that the increase in GFAP-IL6Tg was less pronounced than in aged WT (Figure 4O). Noticeably, in all groups, higher CD68 staining was found in close relationship with microglial clusters, without remarkable differences between genotypes (arrowheads in Figure 4I-L).

d) *MHC-II expression:*

Furthermore, and taking into consideration that FNA lesion induces lymphocyte infiltration in the FN parenchyma (Werner et al. 2001), we analyzed the expression of MHC-II, a molecule importantly related to antigen presentation in the CNS. In basal conditions, MHC-II was strongly expressed in elongated perivascular cells in both aged WT and GFAP-IL6Tg, without differences in their expression or distribution between genotypes (Figure 5C).

**Figure 5.**
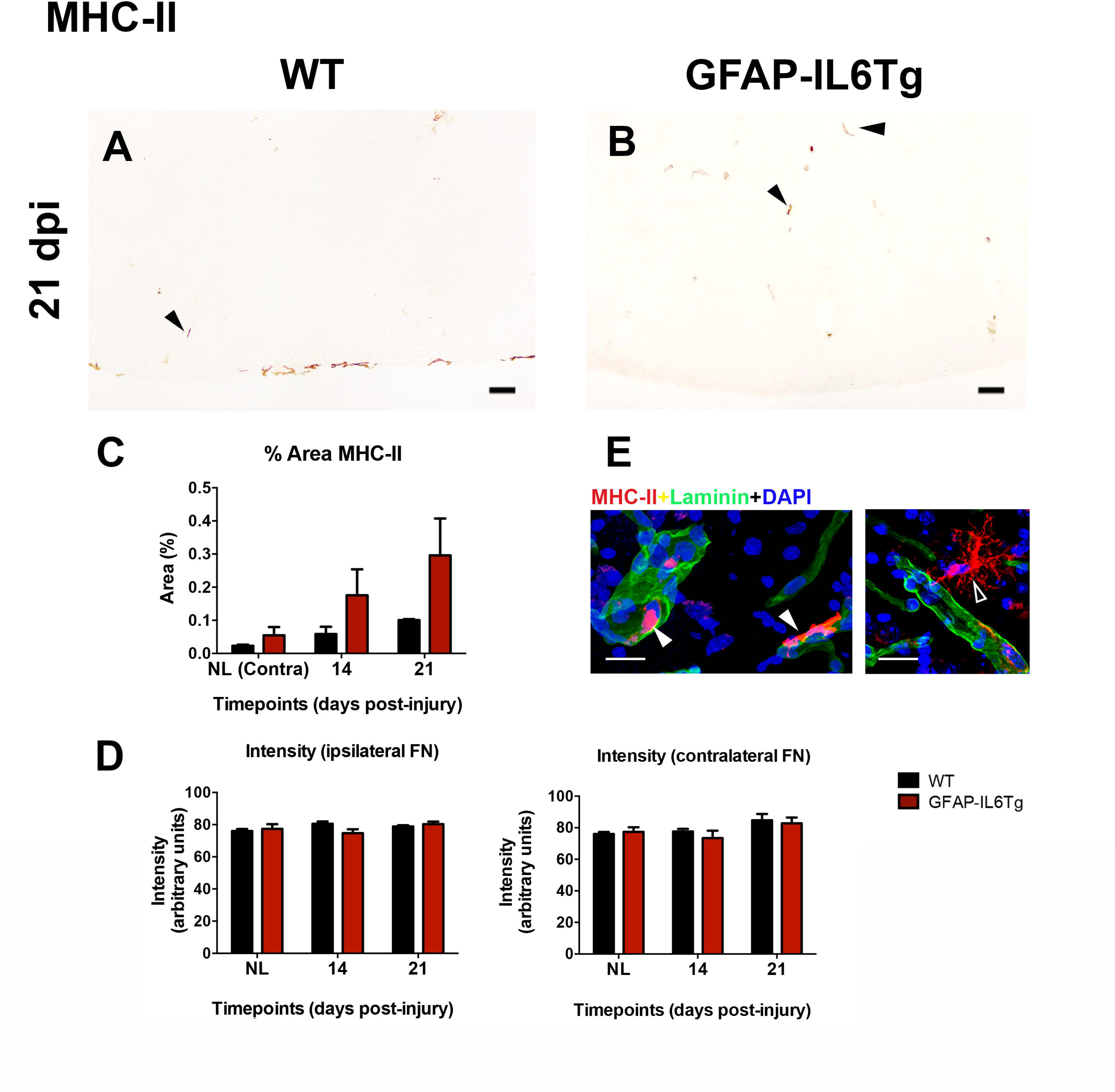
MHC-II expression in basal conditions and after a FNA in aged WT and GFAP-IL6Tg mice. **(A-B)** Representative photomicrographs from MHC-II immunostaining in old WT and GFAP-IL6Tg mice after 21 dpi upon FNA. MHC-II-positive cells are indicated with black arrowheads. **(C)** Quantification of the % Area of MHC-II immunostaining in WT and GFAP-IL6Tg show a tendency to increase at 14 and 21 dpi in WT and GFAP-IL6Tg (2-way ANOVA; time p=0.06 and genotype p<0.05). **(D)** Intensity of MHC-II staining in the ipsilateral and contralateral FN of old WT and GFAP-IL6Tg at all timepoints remains constant. **(E)** Double immunostaining shows MHC-II-positive perivascular macrophages (red, full arrowheads) in contact with the basal lamina (laminin, stained in green) in aged GFAP-IL6Tg mice. MHC-II-positive ramified cells (empty arrowhead) were exceptionally located in the brain parenchyma. Scale bar (A, B)= 50 µm; (E)= 20 µm.

After FNA, a slight increase of MHC-II immunoreactivity was observed in both WT and GFAP-IL6Tg mice. MHC-II expression was found mostly in elongated perivascular cells, and only exceptionally, MHC-II-positive microglia was observed in the brain parenchyma of either aged WT or GFAP-IL6Tg (Figure 5A and B). The quantitative analysis did not demonstrate significant differences at any timepoint in the MHC-II-positive perivascular macrophages between axotomized WT and GFAP-IL6Tg mice in neither the percentage of area nor the intensity of MHC-II (Figure 5A-D).

### Lymphocyte infiltration in basal conditions and after FNA

In addition to modifications in microglia/macrophages, we investigated whether T-lymphocyte infiltration could be altered due to IL-6 overproduction in aged mice, using CD3 as a pan-marker of T-cells. In the contralateral FN of aged NL WT and GFAP-IL6Tg animals a low number of CD3+ cells were found without significant differences among genotypes (Figure 6C).

**Figure 6.**
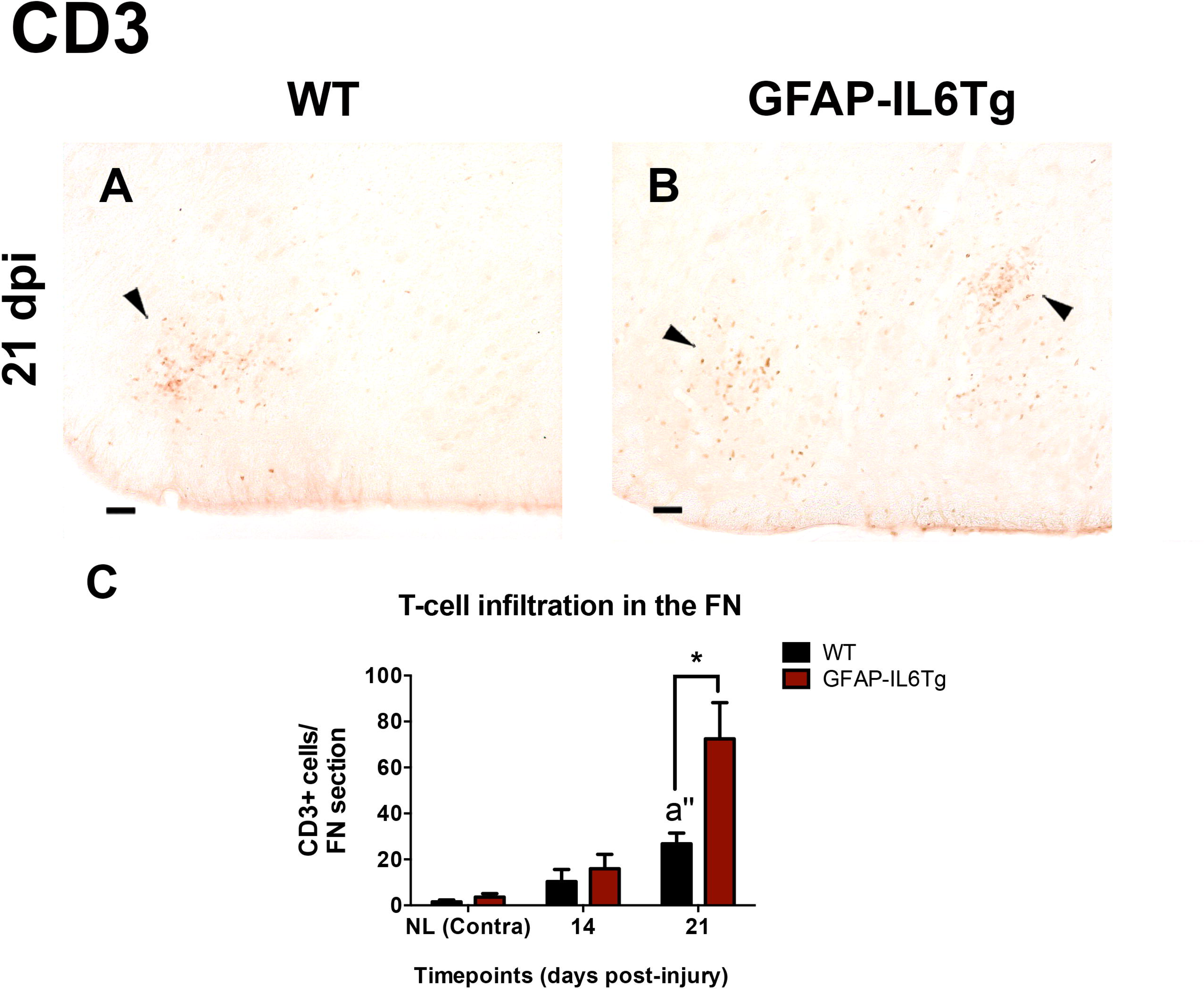
T-lymphocyte infiltration in the FN of aged WT and GFAP-IL6Tg mice after a FNA. **(A-B)** Representative photomicrographs corresponding to the ipsilateral FN of WT and GFAP-IL6Tg at 21 dpi. Infiltrated T-cells are indicated with arrows. **(C)** Quantification of the total number of infiltrated CD3+ cells for each FN section in non-lesioned (NL) animals, contralateral (Contra) and ipsilateral (Ipsi) hemispheres at 14 and 21 dpi in aged axotomized WT and GFAP-IL6Tg mice. An increase in T-lymphocyte infiltration is observed in the lesioned FN at 21 dpi in WT and GFAP-IL6Tg (2-way ANOVA, time ***p<0.001, genotype *p<0.05; post-hoc Tukey’s, a’’ p<0.01), and in the latter, T-lymphocyte infiltration is higher compared to lesioned WT (*p<0.05). Scale bar= 50 μm.

A progressive increase in the number of CD3+ infiltrated T-lymphocytes was detected in the axotomized FN of both genotypes, showing the maximum at 21dpi (Figure 6A-C). Remarkably, a significant increase in the number of CD3+ T-lymphocytes was found in the axotomized GFAP-IL6Tg animals at 21 dpi (Figure 6C).

### Astrocytes in basal conditions and after FNA

Since IL-6 overproduction is generated through the GFAP promoter, putative modifications induced by transgenic IL-6 overproduction in astrocytes were evaluated using GFAP immunostaining. In both aged NL WT and GFAP-IL6Tg mice, GFAP immunostaining showed astrocytes with a star-shaped morphology (Figure 7A, B), some of them showing an intensely stained soma and thick and long projections (Figure 7G and H). The quantitative analysis revealed no differences in either the area or the intensity of GFAP expression between NL aged WT and GFAP-IL6Tg mice (Figure 7K and L).

**Figure 7.**
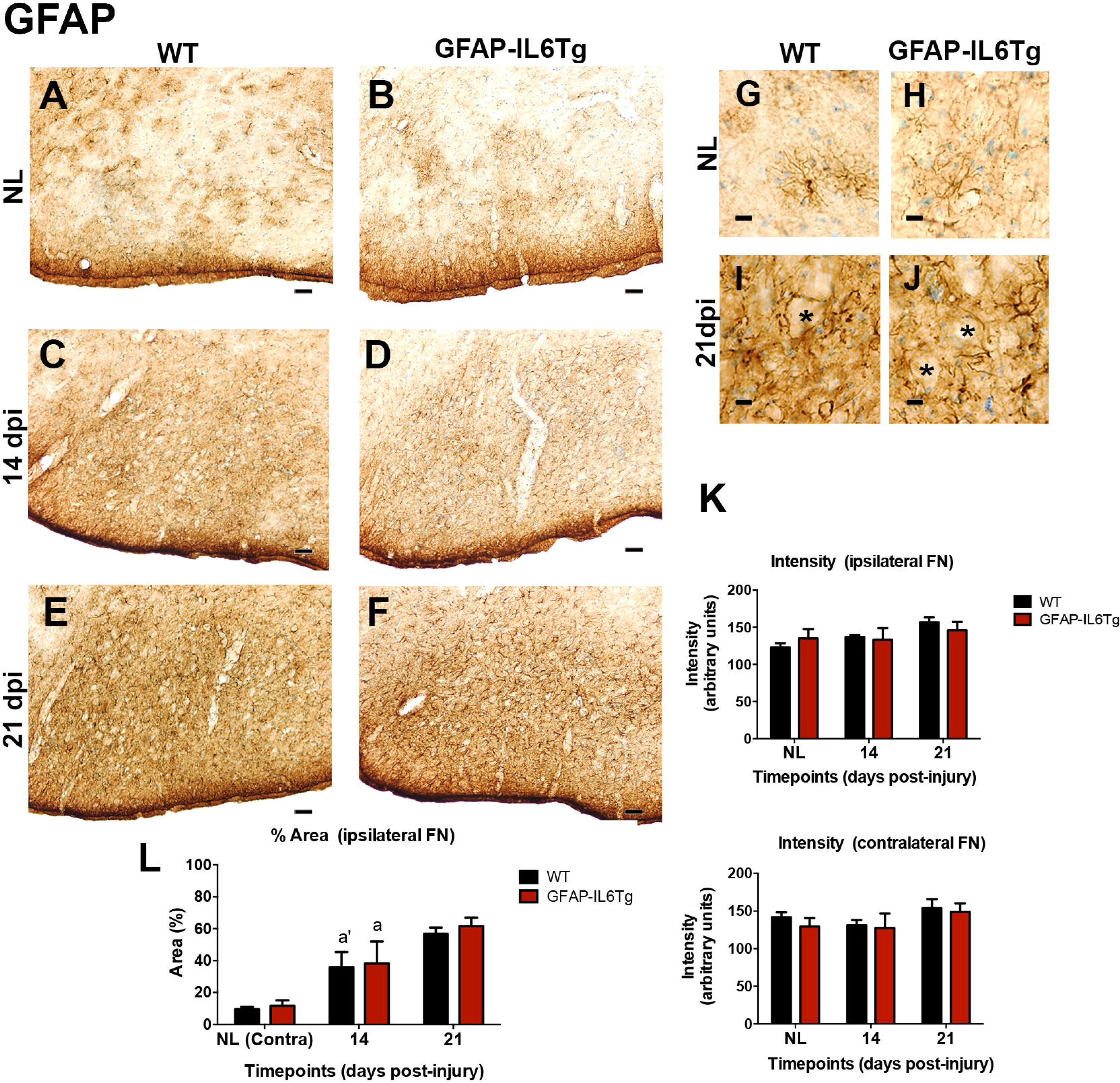
Astroglial reactivity in basal conditions and after FNA in aged WT and GFAP-IL6Tg mice. **(A-F)** Representative photomicrographs from GFAP immunostaining in old WT and GFAP-IL6Tg mice in basal conditions and after peripheral nerve injury. **(G-J)** Magnification showing GFAP+ astrocytes in the FN. Note the hypertrophic morphology in basal conditions in old WT mice (G) and GFAP-IL6Tg (H). After the axotomy, GFAP increases and astroglial reaction is enhanced (I, J), and astrocytes envelop FMN (asterisks). **(K)** Intensity of GFAP staining in the ipsilateral and contralateral FN of old WT and GFAP-IL6Tg at all timepoints does not show statistically significant differences. **(L)** Quantification of the % Area occupied by GFAP in WT and GFAP-IL6Tg reveals similar levels on both mouse lines. Higher levels of % Area are observed at 14 dpi in WT and in GFAP-IL6Tg respect to the previous timepoint (2-way ANOVA, time p<0.0001; Tukey’s multiple comparison, a p=0.05, a’ p<0.05). Scale bar (A-F) = 50 µm. Scale bar (G-J) =10 μm.

After FNA, an increase in GFAP immunostaining was observed (Figure 7C-F) in both genotypes together with morphological changes such as cell hypertrophy (Figure 7I, J). Moreover, GFAP+ projections surrounding FMN were detected, especially at 21dpi in both genotypes (asterisks in Figure 7I, J). The quantitative study showed an increase in the percentage of GFAP+ area at 14 and 21dpi in both WT and GFAP-IL6Tg mice, without differences among genotypes (Figure 7K and L).

### Neuronal survival after FNA in aged WT and aged GFAP-IL6Tg animals

Finally, to determine whether the modifications detected in aged GFAP-IL6Tg in terms of microglial activation and lymphocyte infiltration impact in the response of the lesion, the survival of FMN was analyzed. The total number of FMN was counted in the ipsilateral and contralateral FN of both WT and GFAP-IL6Tg mice using toluidine blue stained sections. In both cases, the contralateral FN of aged WT and aged GFAP-IL6Tg contained around 2000 motor neurons, without significant differences between them (Figure 8E). In both genotypes, the ipsilateral FN of lesioned animals, showed an important reduction of FMN compared to their respective contralateral FN. Thus, in WT, the number of neurons was reduced by 23.85 ± 1.63 % (surviving neurons = 1593 ± 173) while in GFAP-IL6Tg the reduction reached 22.17 ± 2.69 % (surviving neurons = 1491 ± 165). In parallel to the reduction in FMN, the ipsilateral FN of aged WT and GFAP-IL6Tg showed a loss in the cellular organization of their sub-nuclei, and a significant increase in the size of the surviving FMN soma (Figure 8F). No significant differences in either the number of surviving FMN or in their mean diameter were detected between aged WT and aged GFAP-IL6Tg mice (Figure 8E, F).

**Figure 8.**
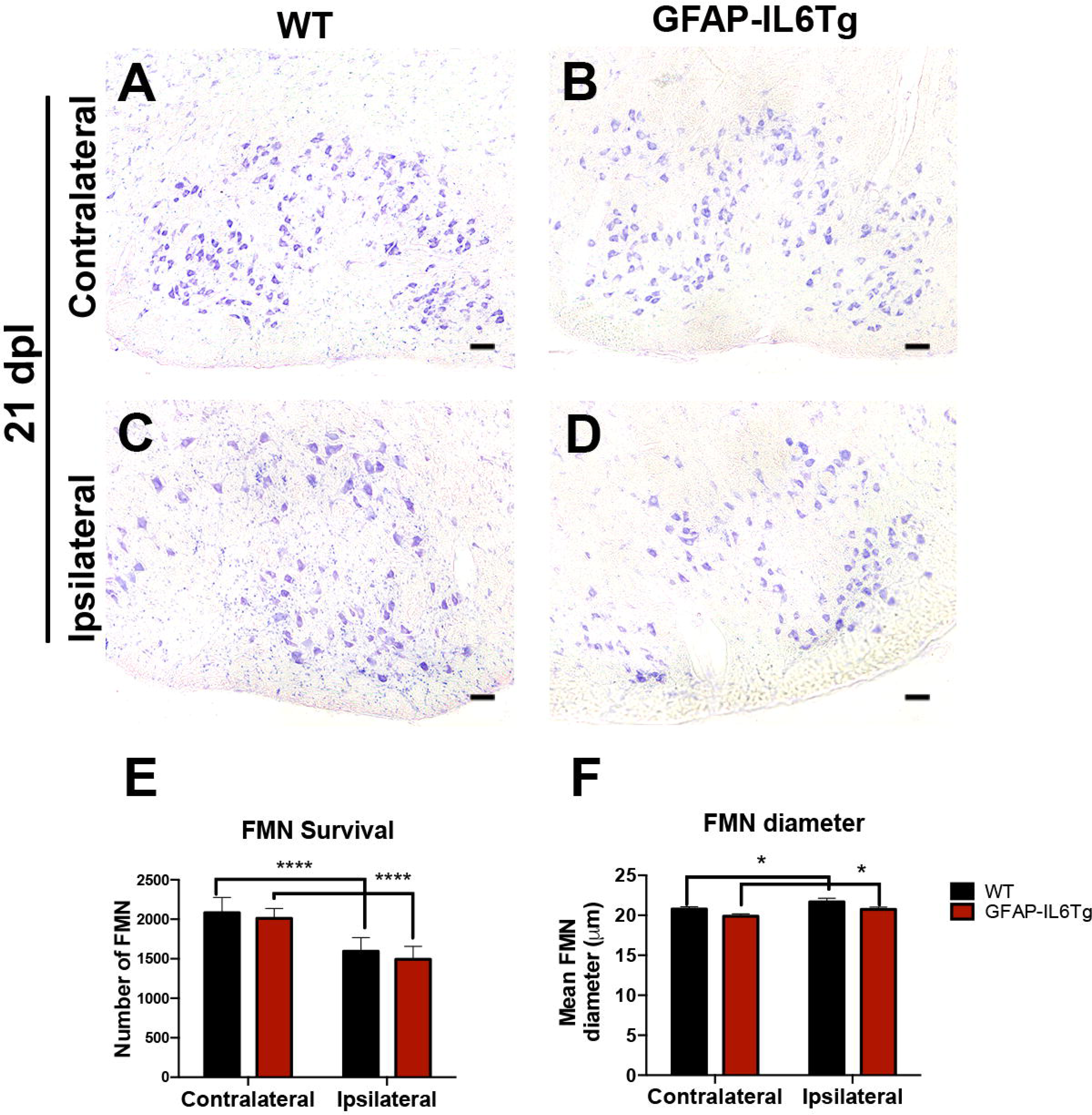
Facial motor neuron survival in aged WT and GFAP-IL6Tg mice after a FNA. **(A-D)** Representative photomicrographs showing toluidine blue stained sections corresponding to the contralateral and ipsilateral FN of old WT and GFAP-IL6Tg at 21 dpi. FMN are easily identified due to the large soma of these cells. **(E)** Quantification of the total number of FMN contained in the whole contralateral and ipsilateral FN of old axotomized WT and GFAP-IL6Tg mice at 21 dpi. Statistically significant reductions between the contralateral and ipsilateral hemispheres of both old WT and GFAP-IL6Tg were found (2-way ANOVA, lesion effect p<0.0001; post-hoc Sidak’s, ****p<0.0001). **(F)** Quantification of the diameter of FMN from lesioned hemisphere showed an increase in the lesioned FMN respect to the non-lesioned (NL) of both old mouse lines (2-way ANOVA, lesion effect p<0.001; genotype effect p=0.06; post-hoc Sidak’s, *p<0.01). Scale bar (A-D) = 50 μm.

## DISCUSSION

Aging is associated with a general pro-inflammatory state, which consistently correlates with increased levels of IL-6, a key pro-inflammatory cytokine upregulated in age-associated neurodegenerative diseases. In the CNS, IL-6 has an overspread regulatory activity in inflammation, with remarkable effects on microglia (Almolda et al. 2014; Recasens et al. 2021; Sanchez-Molina et al. 2021; West et al., 2022a; Rothaug et al., 2016; Spooren et al., 2011; West et al., 2019).

In the present study, we observed that chronic exposure to pro-inflammatory IL-6 induced increased basal microglial reactivity in the FN of aged mice, as clearly detected by higher Iba-1, CD11b, and CD68 microglial expression. These features were compatible with activated microglia described in aging (Henry et al. 2009, Norden and Godbout 2013), and were similar to those previously reported in other brain areas of aged GFAP-IL6Tg mice (Sanchez-Molina et al. 2021). Age-associated microglial activation has been usually linked to primed microglia, in which microglial responds to an inflammatory stimulus in an exacerbated manner. In the last years, the concept of microglial priming has been encompassed in the framework of microglial memory, which includes both trained (or enhanced) and tolerant responses to either priming or desensitizing stimuli, depending on their timing, sequence, strength, or duration (Neher and Cunningham 2019). Previous results from our group showed that chronic IL-6 expression produced microglia activation in NL adult GFAP-IL6Tg mice compared to WT mice, and modified microglia reactivity after CNS lesions, producing either a tolerogenic or an exacerbated response (Almolda et al. 2014; Recasens et al. 2021). These observations led us to describe that microglia under chronic IL-6 expression showed a primed phenotype. Since in this work we found that chronic IL-6 overexpression produced microglia activation in aged mice in basal conditions, our next step was to evaluate how aging shaped the microglial response to chronic IL-6 overproduction upon a CNS lesion. For this purpose, we studied the neuroinflammatory response in aged GFAP-IL6Tg after FNA.

Based on our previous report showing that adult GFAP-IL6Tg mice displayed a primed response to FNA compared to WT mice (Almolda et al. 2014), we expected an exacerbated microglia response in aged lesioned GFAP-IL6Tg compared to their respective WT mice. However, our data showed that, after FNA, chronic IL-6 overproduction in old mice resulted into similar levels of microglial activation markers (Iba1, CD11b, CD68) to that of lesioned aged WT. As a matter of fact, while levels of expression of microglial markers reached similar levels in both mouse lines; aged, lesioned GFAP-IL6Tg animals showed a milder fold-change increase compared to aged, lesioned WT animals. Therefore, our results suggested that, after FNA, microglia of aged GFAP-IL6Tg animals were less activated than aged WT animals. A possible explanation for this observation might be related to a maximum threshold for microglia activation; however, our previous findings in adult lesioned GFAP-IL6Tg mice showed an exacerbated and earlier microglial response compared to adult lesioned WT mice (Almolda et al. 2014). Furthermore, additional evidence from our study supported a milder microglial activation in aged GFAP-IL6Tg. For instance, while in the adult lesioned GFAP-IL6Tg mice the exacerbated microglial response was accompanied by reduced cluster number and FMN survival compared to adult lesioned WT (Almolda et al. 2014), in our study, we did not find differences in FMN survival between aged WT and GFAP-IL6Tg mice nor in microglial cluster number or astrocyte reactivity; and we observed a similar percentage of FMN survival similar to the previously observed in lesioned adult and aged WT mice (Dauer et al. 2011; Almolda et al. 2014). Indeed, the unique parameter that we found altered in aged GFAP-IL6Tg mice was peripheral T-cell infiltration, which was higher in aged, lesioned GFAP-IL6Tg mice compared to WT mice, as similarly observed in lesioned adult transgenic mice after FNA and perforant pathway transection (Almolda et al. 2014; Recasens et al. 2021). These results indicated a putative role of IL-6 in the entrance of immune cells, which could be the result of IL-6 activated endothelial cells, through the increase of ICAM expression, although the involvement of chemokines has also been proposed (Chiang et al., 1994). In addition, in our work we observed an increase of MHC-II expression in aged GFAP-IL6Tg compared to WT mice at 21 dpi, which corresponded mainly to MHC-II-positive perivascular macrophages; and only few microglia expressed MHC-II. Low MHC-II expression in microglia in both aged, lesioned WT and GFAP-IL6Tg suggested a reduced microglial antigen-presentation function, which appeared to be age-specific, since in both adult lesioned WT and GFAP-IL6Tg mice MHC-II expression is more extended in microglia during the peak of microgliosis, principally in clusters (Supplementary Figure 1; Petitto et al., 2003; Villacampa et al., 2015). Remarkably, MHC-II expression in lesioned, aged GFAP-IL6Tg is especially reduced compared to lesioned, adult GFAP-IL6Tg mice (Supplementary Figure 1) suggesting a different effect of IL-6 in microglia from adult or aged mice. Although MHC-II antigen presentation in the microglial compartment has been proved to be essential for FMN survival after FNA (Byram et al., 2003), a putative modulation of the adaptative immune response mediated through the MHC-II downregulation could be reinforced by the lack of T-cell differentiation, as the high number of peripheral infiltrated T-cells in GFAP-IL6Tg may show the similar lack of specification to the T-lymphocyte infiltration observed in the hippocampus of these transgenic animals after perforant pathway transection (Recasens et al. 2021). Our results are opposed to the high levels of MHC-II or CD11b microglial expression associated to an age-related trained microglial phenotype (Henry et al., 2009; Norden et al., 2013). Then, a global analysis of results obtained in this study show that chronic IL-6 overproduction during aging limits microglial response, consequently promoting FMN survival after FNA; and suggests that, within the context of aging and chronic IL-6 overproduction, microglia display a tolerant response to FNA.

The effect of IL-6 in microglial immune memory during aging, and especially after an inflammatory challenge, has been limitedly explored, despite the direct correlation between IL-6 levels and aging. Previous studies demonstrated that IL-6 plays an essential role in microglia priming of aged mice upon peripheral LPS administration (Garner et al. 2018), mainly through the trans-signalling pathway (Burton et al. 2011; Burton and Johnson 2012). In these reports, aged IL-6 knockout animals did not show the increase in IL-1ß, TNF-α, or IL-10 production or upregulated MHC-II microglial expression observed in the hippocampus of aged WT mice after peripheral LPS administration, and these changes were accompanied by the lack of the associated exaggerated sickness behavior observed in LPS-administered aged WT. Our results show an opposite effect, since high levels of IL-6 in aged GFAP-IL6Tg animals produce a tolerant microglial response to FNA, characterized by a lower microglia activation, low MHC-II microglial expression in the brain parenchyma-located out of microglial clusters-, and conserved levels of IL-10 and TNF-α (Supplementary Figure 2). Results related to IL-6 induction of MHC-II expression are contradictory, since IL-6 incubation in BV-2 macrophage cultures stimulated MHC-II expression, while in primary microglia cultures MHC-II was induced only after LPS and IFN-γ, but not IL-6 incubation (Shafer et al., 2002; Garner et al., 2018; Chaufan et al., 2021). Differences in these studies could be explained by both the dose of IL-6 and its incubation with or without its soluble receptor (sIL-6R) (Garner et al., 2018). Since our work is done in vivo, both IL-6 and sIL-6R are expressed, and the effect of IL-6 may be more dependent on the total amount of IL-6 rather than sIL-6R expression, which do not show remarkable differences in levels between aged WT and GFAP-IL6Tg in both NL and lesioned animals (Supplementary Figure 3). Therefore, although the regulation of IL-6R expression have been suggested to be influenced by IL-6 levels (Blanchard et al., 2002), we cannot attribute a diminished microglial response to the downregulation of sIL-6R expression.

Because IL-6 overproduction produced different effects in aged compared to adult mice (tolerant vs. exacerbated responses), we may explain these incongruences through differences in the microenvironment and/or microglial phenotype produced by aging. Specifically, some works showed that IL-6 can have a wide range of effects depending on the specific inflammatory microenvironment in which this cytokine signals, a factor that could modulate the tolerant microglial response to chronic IL-6 overproduction. For instance, IL-6 have a depolarizing effect in primary cultures of microglia stimulated with LPS, shifting microglia from a pro-inflammatory M1 phenotype to a depolarized M0 (Chauhan et al., 2021); but when IL-6 is incubated with either pro-inflammatory (IFN-γ) or anti-inflammatory cytokines (IL-4, IL-13) in cultured macrophages, IL-6 enhances their polarization towards M1 or M2 phenotypes, respectively, suggesting that IL-6 potentiates macrophage commitment (Fernando et al. 2014). In our case, chronic IL-6 overproduction could potentiate an anti-inflammatory or tolerant microglial response within a specific cytokine milieu. Indeed, microglia during aging has been described to show a bias towards an anti-inflammatory phenotype, and to show a sensome more sensitive to internal but not external stimuli (Hickman et al., 2015). Also, tolerant outcomes have been described “in vitro”, in microglia cultures from aged rodents after LPS exposure (Njie et al., 2012; Lajqi et al., 2020). Although, to our knowledge, the specific effect of IL-6 in microglial phenotype upon a central injury has not been studied, several works proved the relevance of cytokine environment related to injury outcome during aging. For instance, a study showed that levels of cytokines upon LPS administration (consisting of IL-1α, IL-2, IL-4, IL-10, and TNF-α) in the brainstem of young rats closely mimicked those of aged rats in basal conditions (Katharesan et al. 2016), and that LPS administered in adult rats before facial nerve avulsion reduced FMN death (Katharesan et al. 2018). In view of our results and the current literature, it may be interesting to deepen in the effect of the cytokine environment in the modulation of the IL-6 effect during aging.

## CONCLUSIONS

In this study, we determined that astrocyte-targeted IL-6 overproduction in aged mice induces an activated microglial phenotype indicative of priming under baseline conditions. After FNA, chronic IL-6 overproduction in aged mice induces a microglial response with protective or tolerant features, that does not affect astroglial activation, and which avoids increased FMN from death, as contrarily observed in adult transgenic GFAP-IL6Tg animals. In view of the conclusions reported in this work, it may be of interest to deepen in the role of IL-6 in microglial priming and tolerance during aging, and its consequences on CNS lesions.

## Acknowledgments

The authors would like to thank Miguel A. Martil and Isabella Appiah for their outstanding technical help. This work was supported by the Spanish Ministry of Science and Innovation (BFU2017-87843-R) to BCL.

## CRediT authorship contribution statement

### Gemma Manich

Writing-original draft, Visualization, Validation, Supervision, Formal analysis, Conceptualization. **Ruggero Barbanti**: Writing-original draft, Validation, Visualization, Methodology, Formal analysis, Data curation. **Marta Peris:** Validation, Methodology, Formal analysis, Data curation. **Nadia Villacampa**: Writing-review and editing, Visualization, Validation, Formal analysis, Data curation. Beatriz Almolda: Writing-review and editing, Visualization, Validation, Conceptualization. **Berta González**: Writing-review and editing, Conceptualization, Project administration, Visualization, Supervision. **Bernardo Castellano**: Conceptualization, Visualization, Project administration, Resources, Funding acquisition.

**Suppl. Figure 1. MHC-II expression in adult WT and GFAP-IL6Tg mice after FNA. (A-B)** Representative microphotographs corresponding to the ipsilateral side of the FN in WT and GFAP-IL6Tg mice at 14 dpi. **(C-D)** High magnification photographs showing the characteristic ramified morphology of MHC-II+ cells found in WT and GFAP-IL6Tg mice. **(E-L)** Triple immunohistochemistry combining Iba1 (green), MHC-II (red) and CD3 (blue). Cell nuclei were stained with DAPI (white). Merged images (H and L) showed colocalization of MHC-II and Iba-1. Arrows point to an Iba1+ microglial cluster in WT (E-H) and GFAP-IL6Tg mice (I-J) containing MHC-II+ cells (F and J). Note that CD3+ cells are close to the microglial cluster (G and K). Scale bar (A and B)= 50µm; (Squares in A and B)= 10µm; (C-J)= 25µm.

**Suppl. Figure 2. IL-10 and TNF-a expression in the FN of aged WT and GFAP-IL6Tg mice during basal conditions and after FNA.** Quantification of IL-10 and TNF-expression was performed in the NL (non-lesioned) and lesioned (21 dpi) FN of aged WT and aged GFAP-IL6Tg. No differences statistical differences were observed (2-way ANOVA, no effect in time and genotype; post-Tukey comparison’s test, no significant differences).

**Suppl. Figure 3. IL-6R expression in the FN of aged WT and GFAP-IL6Tg mice during basal conditions and after FNA.** Quantification of IL-6R was performed in the NL (non-lesioned) and lesioned (21 dpi) FN of aged WT and aged GFAP-IL6Tg. Statistical differences showed an effect of lesion (2-way ANOVA, lesion ** p<0.001 and genotype n.s.; post-Tukey comparison’s test, n.s.).

