## Supplementary figures and images for "Chronic IL-6 overproduction induces a tolerogenic response in aged mice after peripheral nerve injury"

### Supplementary Figure 1

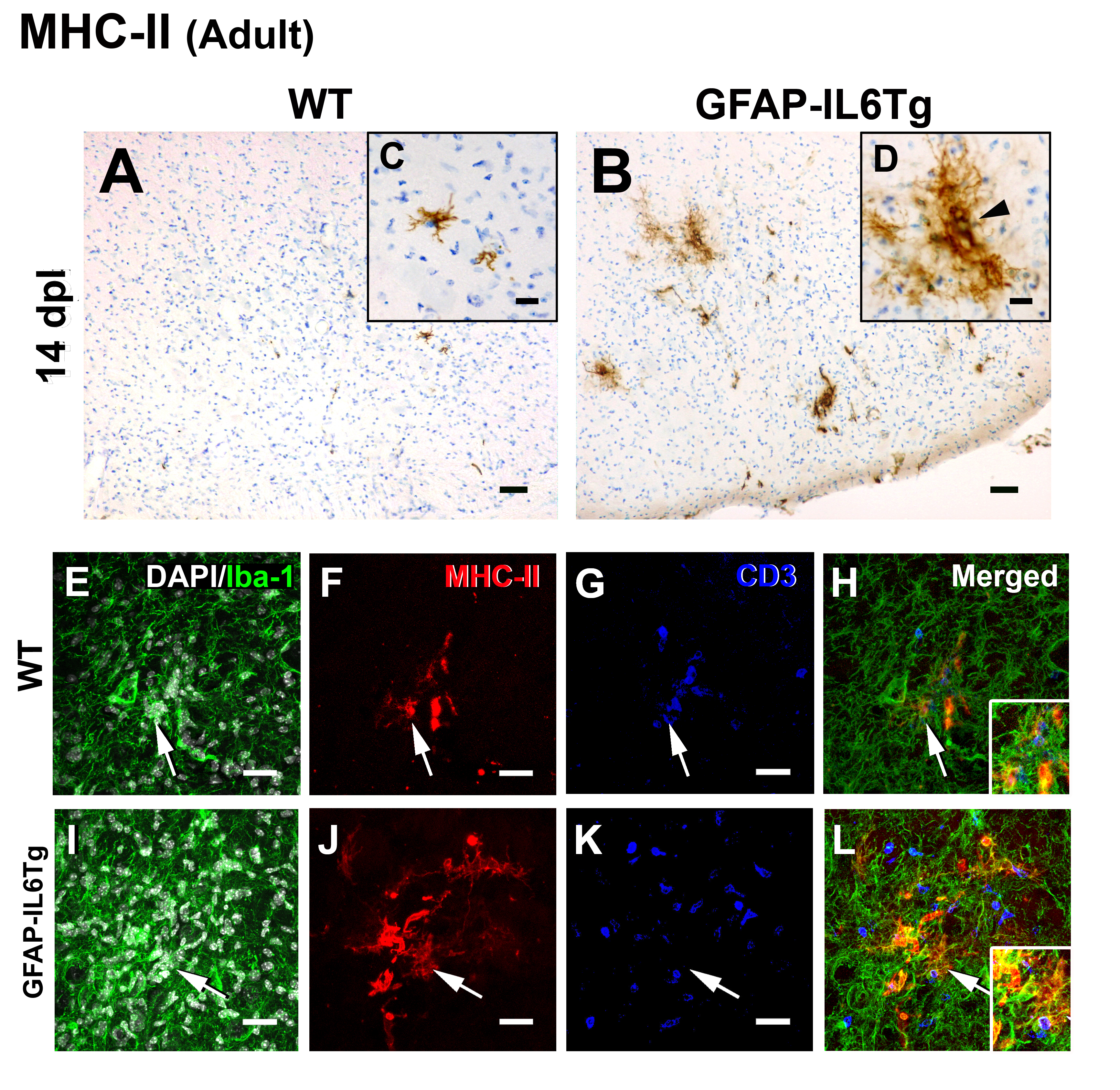

### Supplementary Figure 2

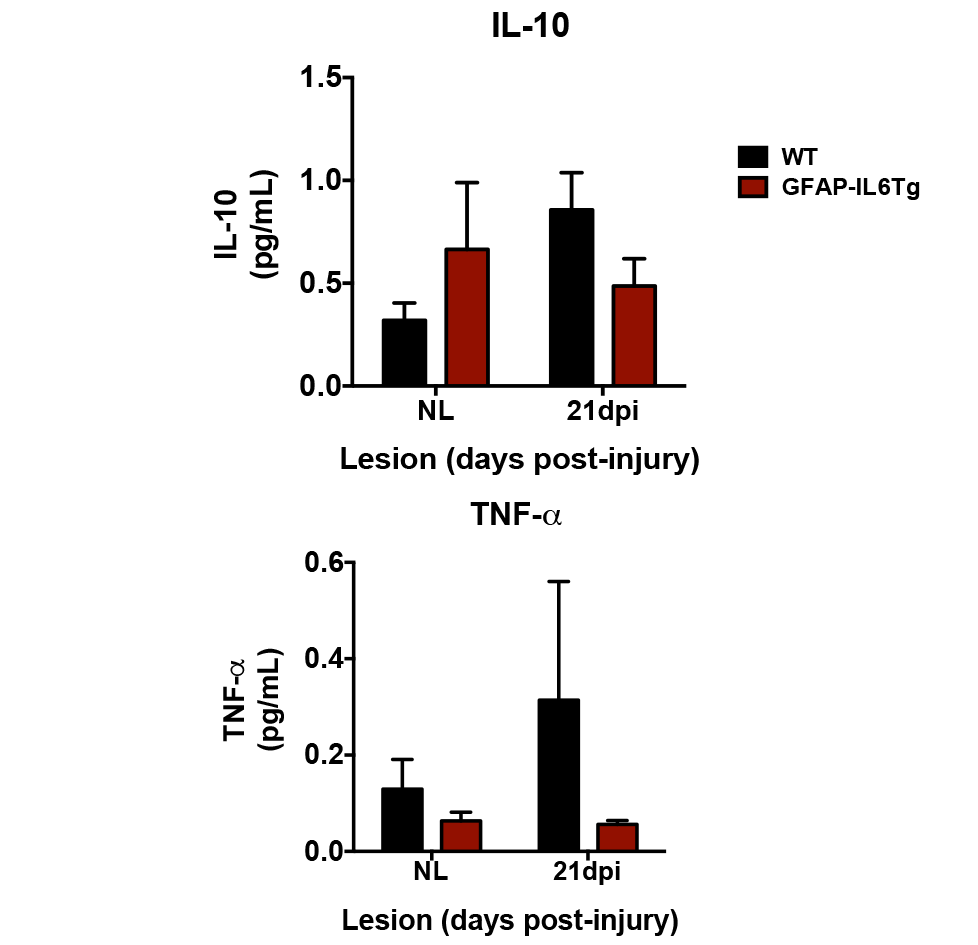

### Supplementary Figure 3

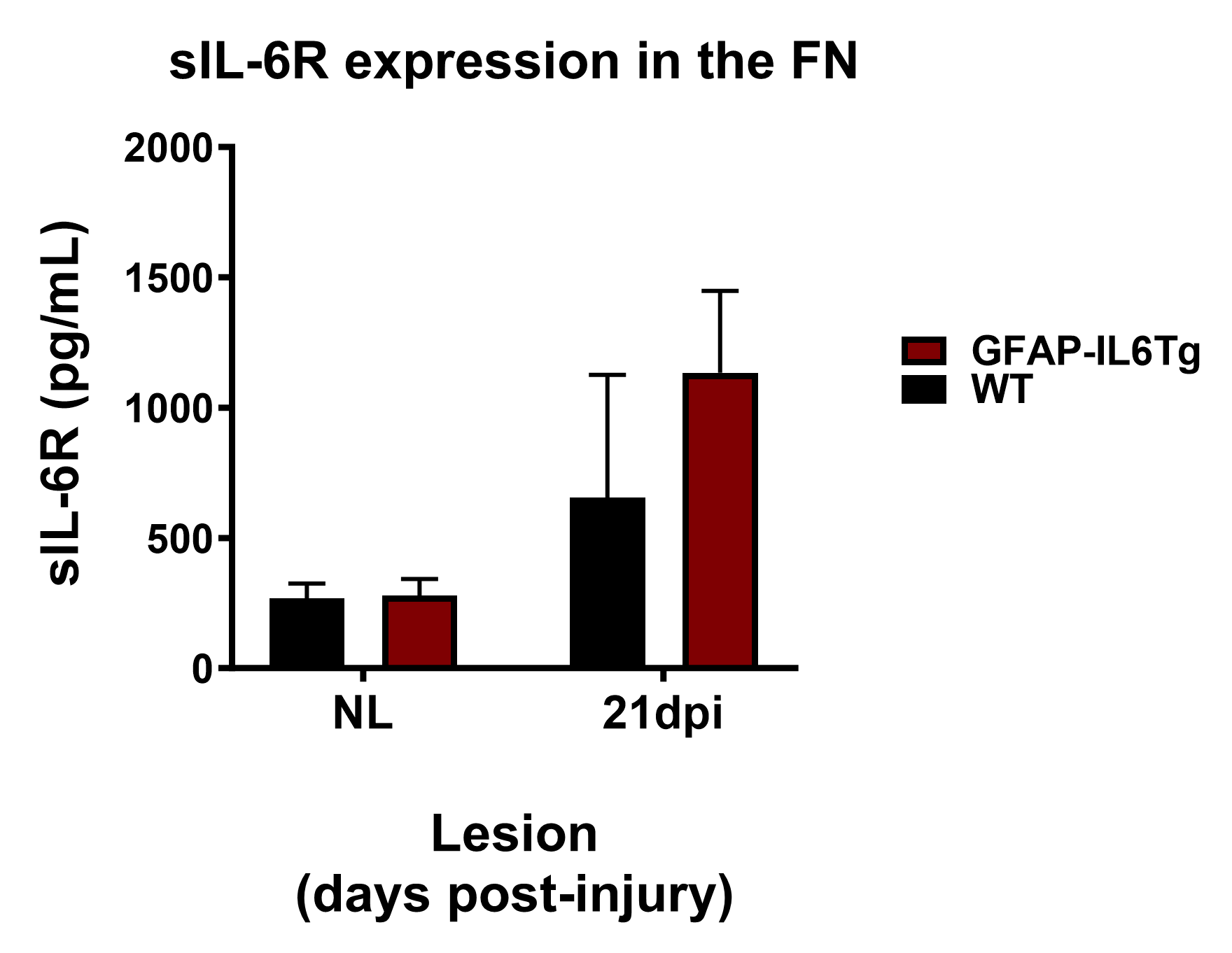
